# An automated metabolite extraction workflow for global metabolomics analysis using the Agilent Bravo liquid handling platform

**DOI:** 10.1101/2025.04.07.647543

**Authors:** Fraser Massie, Andrew MacKenzie, Sufyan Pandor, Mauro A. Cremonini, Tessa Moses

**Author notes:** These authors contributed equally to this work (Fraser Massie, Andrew MacKenzie).

## Abstract

Metabolomics is the comprehensive study of small molecules that provides a snapshot of an organism’s physiological state. Reflecting phenotype more closely than genes or proteins, metabolites reveal changes linked to diseases, mutations, genetic interventions, and environmental stimuli. Recent technological advancements in metabolomics analysis through the application of ion mobility mass spectrometry have enhanced the comprehensive analysis of complex metabolic mixtures. However, pre-analytical bottlenecks in throughput and consistent extraction persist. We developed an automated, sample-agnostic metabolite extraction workflow for diverse liquid samples using an Agilent Bravo liquid handling platform. Here, we provide device, protocol, and form files for efficient sample processing to extract metabolites for global metabolomics analysis using liquid chromatography – mass spectrometry techniques.

## Introduction

Metabolites are small molecules with low molecular weight, typically under 1,500 Da, that are the end products of cellular metabolism. They include a wide variety of chemical groups, including sugars, amino acids, lipids, organic acids, and various other compounds. Metabolomics, often referred to as global or untargeted metabolomics, is the comprehensive study of these metabolites within a biological system (Patti et al., 2012; Trivedi et al., 2017). Metabolomics aims to detect, identify and quantify all the metabolites present in a sample, thus providing a snapshot of an organism’s physiological state at the time of sampling.

Metabolites are a direct reflection of an organism’s phenotype (Fiehn, 2002). Small molecule metabolites are closer to the observable characteristics (phenotype) than genes or proteins. Changes in metabolite levels can directly reflect physiological changes, diseases, or responses to external stimuli, thus making their analysis through metabolomics important for understanding biology (Nicholson et al., 1999). Metabolomics offers a systems-level view of biological processes, complementing genomics and proteomics (Resurreccion and Fong, 2022; Akyol et al., 2023). It helps understand interactions between different metabolic pathways and their overall impact on the organism. By analysing metabolic profiles, we can gain insights into the regulation and function of metabolic pathways, thus bettering our understanding of cellular processes, which can be used for applications such as disease prognosis and treatment. Metabolomics can also be used to identify biomarkers of disease, responses to drug treatments, and environmental exposures, which has implications for diagnosis, personalised medicine and monitoring treatment efficacy (Collino et al., 2013; Ford et al., 2022).

Recent advancements in metabolomics instrumentation, coupled with powerful analytical techniques like ion mobility mass spectrometry (IMMS), have enabled more comprehensive analysis of complex biological mixtures (Moses and Burgess, 2023). IMMS integrates the separation power of ion mobility (IM) with the identification capabilities of mass spectrometry (MS). Furthermore, rapid IMMS-based methods have been developed for high-throughput metabolomics analysis (Pičmanová et al., 2022), significantly reducing analytical time and increasing sample throughput, allowing for the processing of hundreds of samples daily. Despite these advancements in analytical capabilities, a bottleneck persists in the pre-analytical stage, specifically the lack of robust, high-throughput sample extraction workflows. To address this limitation, we developed an automated, sample-agnostic, multi-sample metabolite extraction workflow suitable for a wide range of liquid samples, including culture media, urine, plasma, serum, cerebrospinal fluid, tears, nasal discharge, saliva, beverages, and wastewater, amongst others (Figure 1). Here, we present the Agilent Bravo liquid handling platform configuration and provide the necessary device, protocol, and form files to expedite metabolite extraction from liquid samples.

**Figure 1.**
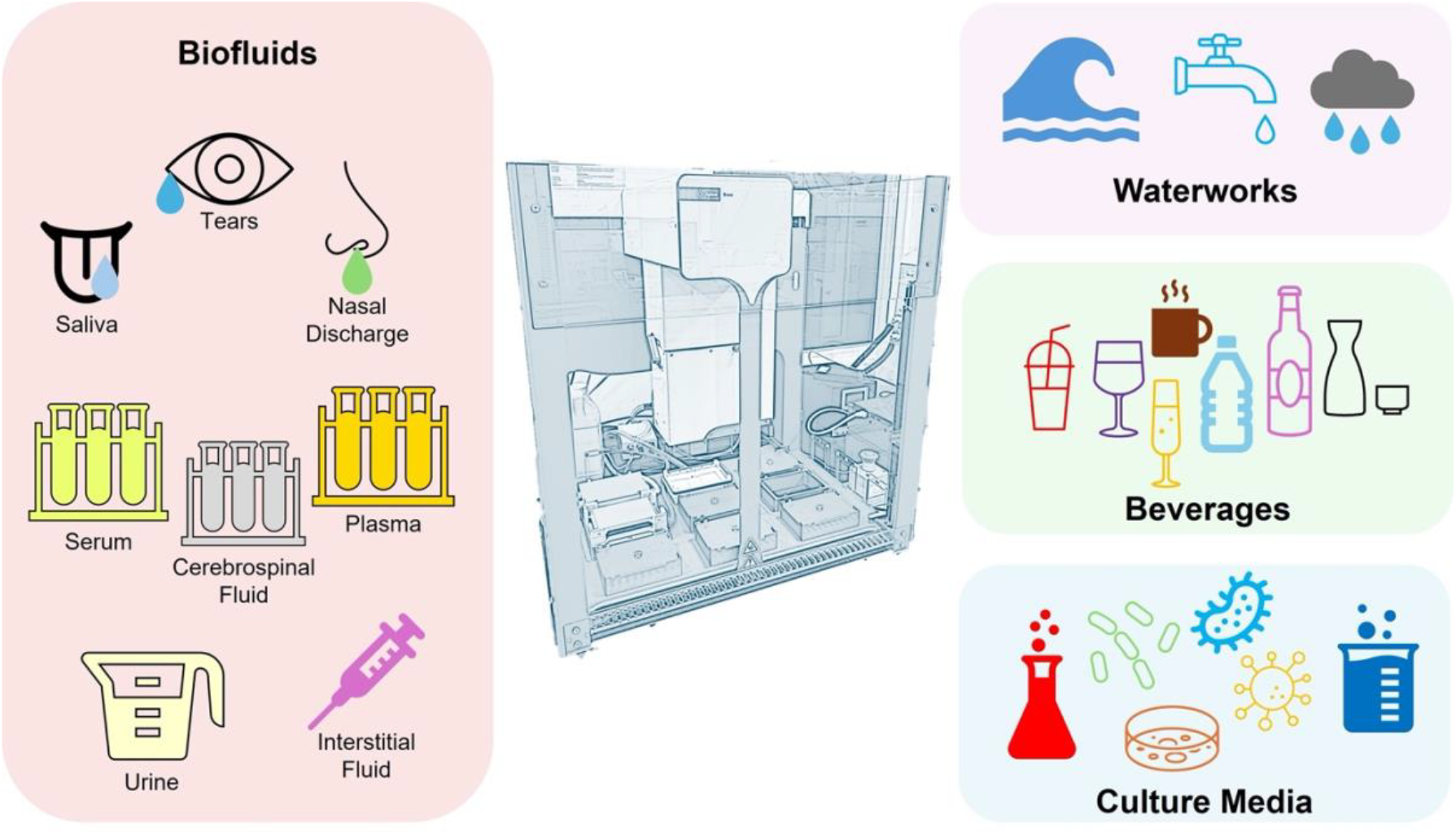
A schematic representation of the Agilent Bravo liquid handling platform and a wide range of liquid samples that can be extracted.

### Experimental

#### Materials

Bravo automated liquid handling platform (Agilent G5563AA) with 96LT 96-well Agilent disposable-tip pipette head (G5055A/G), heating shaking station (Agilent G5498B/G #009), recirculating heater/chiller (Agilent G5498B/G #024), thermal station (short-side connectors; Agilent G5498B/G #036), vacuum filtration station with pump (Agilent G5432B/G) and 1.5 mL microtube rack adapter (Agilent G5477-20006). Reservoir, single cavity, polypropylene, 86 mL, 96 pyramids base geometry, 19 mm height, 25/pk (Agilent 201254-100) for holding extraction solvent. Tips, 250 µL, 96 in rack, case of 50 compatible with Bravo 96LT head (Agilent 19477-002). Microcentrifuge tube, natural polypropylene, 1.5 mL capacity. Temperature-controlled bench top centrifuge with microcentrifuge tube rotor. Fisherbrand™ 11 mm plastic snap ring micro-vial with glass micro-insert (11585914) and SureSTART™ 11 mm snap caps (Thermo Scientific 6PRC11ST1) for high-performance liquid chromatography (HPLC) applications. In-house designed and 3D printed 24 well LC vial adapter rack (.step file can be accessed on GitHub).

#### Agilent Bravo configuration

The Agilent Bravo deck can be customised for user-specific needs. The workflows discussed here are written for a Bravo platform configured with a cooling unit on position 1, a vacuum filtration station on position 2, and a heating shaker station on position 4. These positions are fixed but compatible with labware. Positions 3, 5, 6, 7, 8 and 9 are deck spaces compatible with various labware.

#### VWorks software setup

1. Before running any workflows, ensure that the protocol files are linked to the ‘96LT w Vac Flt Stn_v4.dev’ device file and the correct form file (Table 1). **NOTE:** In the protocol file navigate to the ‘Protocol Options’ tab, and under ‘Properties’, specify the device and form files using the ‘Device file path’ and ‘Form to use’.
2. Ensure that the ‘Start’ button on each form file links to the correct protocol file. In VWorks software select ‘Edit Form’ in the ‘Tools’ drop-down menu, then click on the ‘Start’ button element in the form editing window. In the ‘Run Specified Protocol Properties’ pane on the right-hand side of the window, ensure that the ‘Protocol file’ field leads to the correct protocol file as detailed previously, then click ‘OK’ and save the form file in VWorks.

**Table 1.**
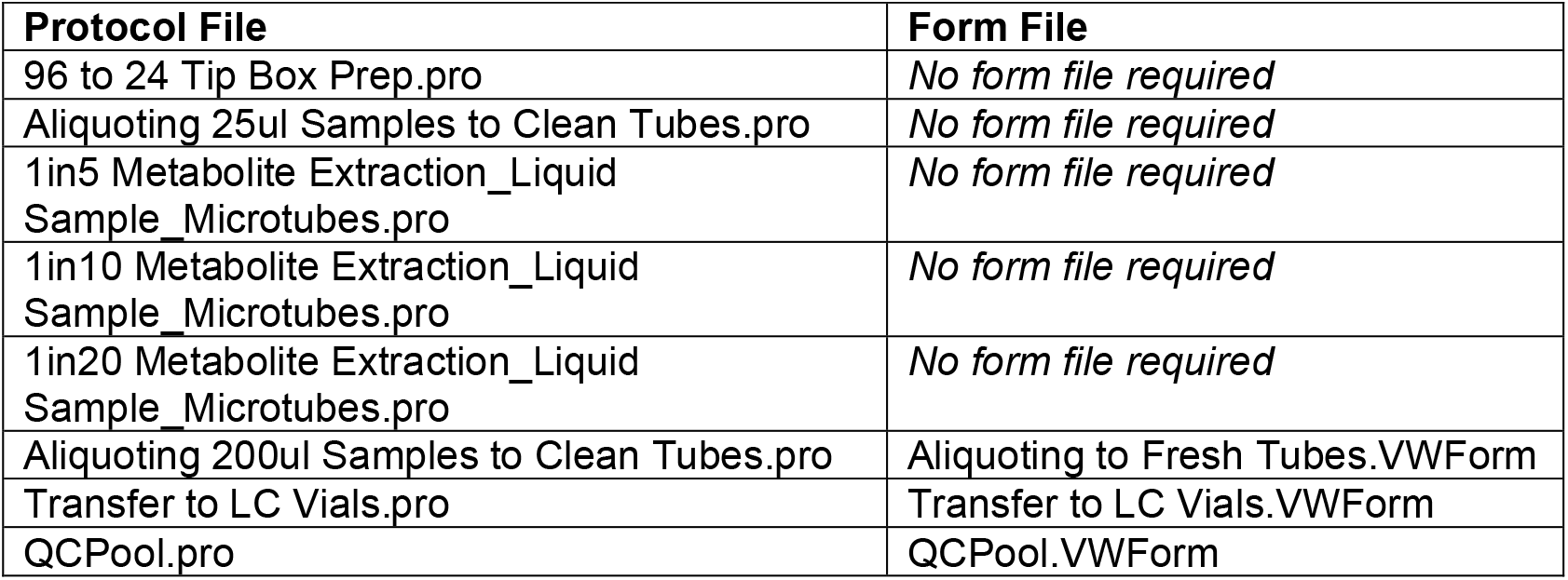
Protocol and corresponding form files to be linked.

### Method files

All device, protocol, form and step files can be accessed on GitHub under different branches
https://github.com/EdinOmics/Automated-metabolite-extraction-workflow-on-Agilent-Bravo-liquid-handling-platform/branches

### Workflows

#### Preparing tip boxes for metabolite extraction of samples in 1.5 mL microcentrifuge tubes

1. Start the Agilent Bravo automated liquid handling platform, the heating shaking station, and the vacuum filtration station pump.
2. Open the VWorks software and load the ‘96 to 24 Tip Box Prep.pro’ protocol (access on GitHub). A prompt will appear to initialise the device now, if the device has not already been initialised. Click ‘Yes’ if applicable.
3. A dialogue box appears stating ‘A plate is present in or in front of plate presence sensor’. Choose ‘Ignore and Continue, leaving device in current state’. A second dialogue box appears stating ‘Please verify it is safe to home the W-axis’. Choose ‘Retry’.
4. Place a full tip box with 96 tips at position 5, and clean empty tip boxes at positions 1, 3, and 6 of the Bravo deck (Figure 2). Use the deck image in the protocol file’s ‘Task Parameters’ section as a positional guide.
5. Click ‘Start’ on VWorks software. A run configuration wizard opens with four pages: (1) Run protocol this many times; (2) Determine when the protocol will start; (3) Select the starting barcode for each; (4) Add notes about the run. Leave all parameters at default settings and click ‘Finish’.
6. A dialog box will open to confirm labware placement. Click ‘Continue’, if the image matches your actual deck placement of parts. A second user message opens to confirm where the full tip box and empty tip boxes are placed. Click ‘Continue’, if all parts are correct on the Bravo deck.
7. The robot will then distribute the tips evenly between the four tip boxes, resulting in 24 tips per box that align with the microtube rack adapter and the 24 well LC vial adapter rack.

**Figure 2.**
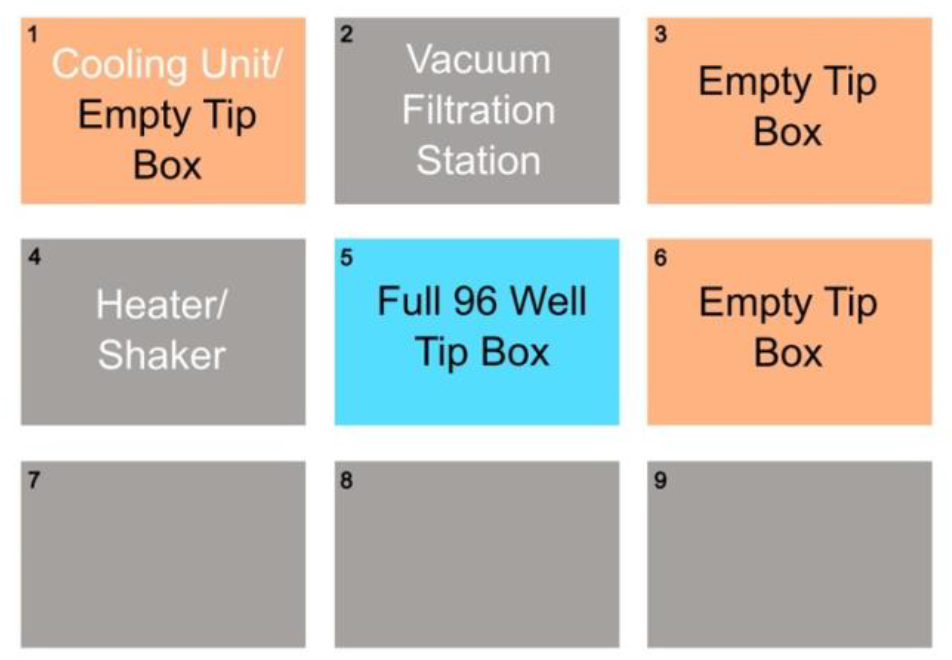
Map of the Bravo deck with labware configured for the ‘96 to 24 Tip Box Prep.pro’ protocol. Fixed labware (white text), full 96 tip box (blue), empty tip boxes (orange), and positions with no movable labware (grey).

#### Aliquoting samples into 1.5 mL microcentrifuge tubes for metabolite extraction

1. Start the Agilent Bravo automated liquid handling platform, the heating shaking station, and the vacuum filtration station pump.
2. On the VWorks software, load the ‘Aliquoting 25ul Samples to Clean Tubes.pro’ protocol (access on GitHub). A prompt will appear to initialise the device now, if the device has not already been initialised. Click ‘Yes’ if applicable.
3. Use the deck image in the protocol file’s ‘Task Parameters’ section to prepare the Bravo deck (Figure 3).
4. Fill a 24-well microtube rack adapter with the 1.5 mL microcentrifuge tubes containing samples. Ensure lids are open and secured in place on the rack. Place this rack on position 4 of the Bravo deck.
5. Fill a second 24-well microtube rack adapter with clean labelled 1.5 mL microcentrifuge tubes. Ensure the tube lids are open and place this rack on position 5 of the Bravo deck.
6. Place a tip box configured with 24 tips on position 8.
7. Click ‘Start’ on VWorks software. A run configuration wizard opens with four pages: (1) Run protocol this many times; (2) Determine when the protocol will start; (3) Select the starting barcode for each; (4) Add notes about the run. Leave all at default settings and click ‘Finish’.
8. The bravo pipette head will pick up all 24 tips from position 8 and move to position 4 to aspirate 25 µL from the sample tubes. **NOTE:** post sample aspiration, a 10 µL volume of air is aspirated into the tip to avoid sample dripping and loss during the transfer process.
9. The tips with the sample then move to position 5, and dispense the aliquoted samples into the clean microcentrifuge tubes.
10. Finally, the tips are returned to position 8, where they are deposited in the tip rack for disposal.

**Figure 3.**
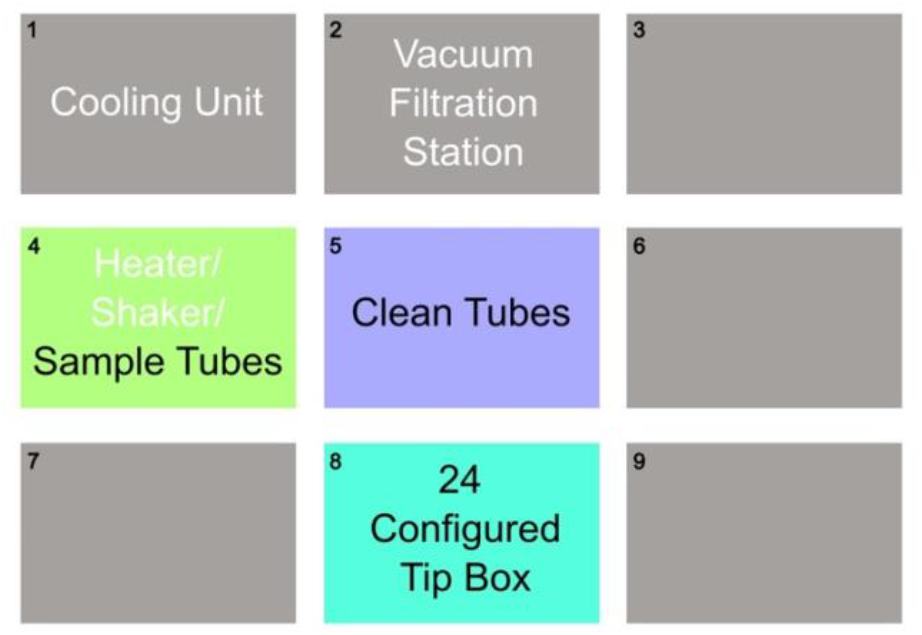
Map of the Bravo deck with labware configured for the ‘Aliquoting 25ul Samples to Clean Tubes.pro’ protocol. Fixed labware (white text), 24-well microtube rack adapter with sample tubes (green), 24-well microtube rack adapter with clean tubes (purple), 24 tip box (turquoise), and positions with no movable labware (grey).

#### Metabolite extraction in 1.5 mL microcentrifuge tubes

1. Start the Agilent Bravo automated liquid handling platform, the heating shaking station, and the vacuum filtration station pump.
2. Switch on the recirculating heater/chiller unit and allow the thermal station at position 1 of the Bravo deck to cool to 4°C.
3. Place a reservoir on position 1 and fill with extraction solvent (Optima MS grade chloroform: methanol: water, 1:3:1, in our case), the microtube rack adapter with 25 µL of sample to be extracted on position 4 (ensure lids are open) and a tip box configured with 24 tips on position 8 (Figure 4).
4. Load the protocol file ‘1in5 Metabolite Extraction_Liquid Sample_Microtubes.pro’, ‘1in10 Metabolite Extraction_Liquid Sample_Microtubes.pro’, or ‘1in20 Metabolite Extraction_Liquid Sample_Microtubes.pro’ (all VWork protocols can be accessed on GitHub). A prompt will appear to initialise the device now, if the device has not already been initialised. Click ‘Yes’ if applicable. **NOTE:** The protocol file to use is determined by the ratio of sample to extraction solvent required for the given experiment. For instance, if a ratio of 1:5 sample: extraction solvent is desired (i.e. 25 µL sample and 100 µL extraction solvent), use the protocol ‘1in5 Metabolite Extraction_Liquid Sample_Microtubes.pro’.
5. Click ‘Start’ on VWorks software. A run configuration wizard opens with four pages: (1) Run protocol this many times; (2) Determine when the protocol will start; (3) Select the starting barcode for each; (4) Add notes about the run. Leave all parameters on default settings and click ‘Finish’.
6. The Bravo head will first pick up all 24 tips from position 8, move to position 1 and aspirate the appropriate volume of extraction solvent from the reservoir into each tip. **NOTE:** post sample aspiration, a 10 µL volume of air is aspirated into the tip to avoid extraction solvent dripping and loss during the transfer process.
7. The tips then move to position 4 and dispense the extraction solvent into the sample tubes. The sample and extraction solvent mixture is mixed for 15 min at 1,200 rpm. For 1:10 and 1:20 ratio of sample: extraction solvent, the solvent is dispensed in two steps and step 7 is repeated. For 1:10 ratio on 25 µL sample, 100 µL extraction solvent is dispensed first followed by 125 µL extraction solvent. And for 1:20 ratio on 25 µL sample, 237.5 µL extraction solvent is dispensed twice. **NOTE:** between the two dispensing steps, the tips are returned to the original tip box for re-use. Following the second dispensing, the empty tips are returned to position 8 for re-use in step 12.
8. The protocol ends after the mixing finishes.
9. The operator can now remove the microtube rack adapter from the deck, close the 1.5 mL microcentrifuge tubes and transfer them to a temperature-controlled benchtop centrifuge for centrifugation at 4°C for 1 min at 21,130 ×g. **NOTE:** This step ensures the pelleting and removal of macromolecules, such as proteins, that might have precipitated during metabolite extraction. The resulting supernatant corresponds to the metabolite extract that will be retained for LC-MS analysis.
10. After centrifugation, the microcentrifuge tubes are carefully placed in the microtube rack adapter with open lids and returned to position 4 of the Bravo deck. A clean set of labelled 1.5 mL microcentrifuge tubes are placed with open lids in a microtube rack adapter on position 5. The tips used in step 7 remain at position 8 (Figure 3).
11. In VWorks software load the protocol file ‘Aliquoting 200ul Samples to Clean Tubes.pro’ (access on GitHub). The form file ‘Aliquoting from Fresh Tubes.VWForm’ (access on GitHub) will then open automatically. Choose the desired volume of solvent to be transferred to the fresh tubes from the drop-down menu. Then click the start button. **NOTE:** Choose 90 µL, 200 µL and 400 µL for the 1:5, 1:10 and 1:20 extraction ratio, respectively.
12. The Bravo head will pick up all 24 tips from position 8, move to position 4 and aspirate the specified volume from the sample tubes. **NOTE:** post sample aspiration, a 10 µL volume of air is aspirated into the tip to avoid extraction solvent dripping and loss during the transfer process.
13. The tips then move to position 5 and dispense the metabolite extract into fresh microcentrifuge tubes, in which they will be held for long-term storage in an ultra-low temperature freezer.
14. Finally, the tips are returned to position 8, where they are deposited in the tip rack for disposal.
15. The original sample extraction microcentrifuge tubes at position 4, containing the residual pellet, can also be disposed at the end of this workflow.

**Figure 4.**
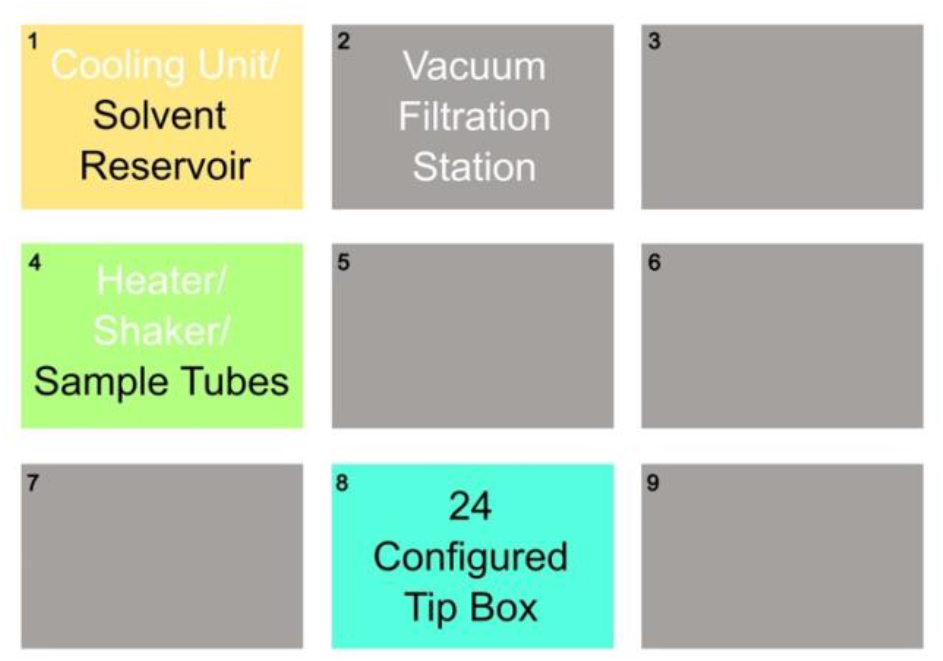
Map of the Bravo deck with labware configured for liquid metabolite extraction protocols. Fixed labware (white text), solvent reservoir (yellow), 24-well microtube rack adapter with sample tubes (green), 24 tip box (turquoise), and positions with no movable labware (grey).

#### Aliquoting metabolite extract into LC vials

1. Start the Agilent Bravo automated liquid handling platform, the heating shaking station, and the vacuum filtration station pump.
2. On the VWorks software, load the ‘Transfer to LC Vials.pro’ (access on GitHub). A prompt will appear to initialise the device now, if the device has not already been initialised. Click ‘Yes’ if applicable.
3. The form file ‘Transfer to LC Vials.VWForm’ will open automatically.
4. Retrieve metabolite extracts from ultra-low temperature freezer and centrifuge at 21,130 ×g for 5 min at 4°C.
5. Place the microcentrifuge tubes with lids open in the microtube rack adapter on position 4 of the Bravo deck.
6. Place the 24 well LC vial adapter rack with labelled LC vials on position 5 of the Bravo deck. Ensure the LC vials are at the analogous position as the sample. Place a tip box configured with 24 tips on position 8 (Figure 5). **NOTE:** This LC vial adapter rack is not available for purchase and was designed and 3D printed in-house (see rack dimensions in Figure 6 and the .step file can be accessed on GitHub).
7. On the form file choose the desired volume of solvent to be transferred to the LC vials from the drop-down menu. Then click the start button. **NOTE:** Typically, 50 µL of the sample extract is transferred to the LC vial.
8. The Bravo head will pick up all 24 tips from position 8, move to position 4, aspirate the specified volume from the sample microcentrifuge tubes, move to position 5, and dispense the extracts into the LC vials. The tips are then returned to the tip rack on position 8 for disposal.

**Figure 5.**
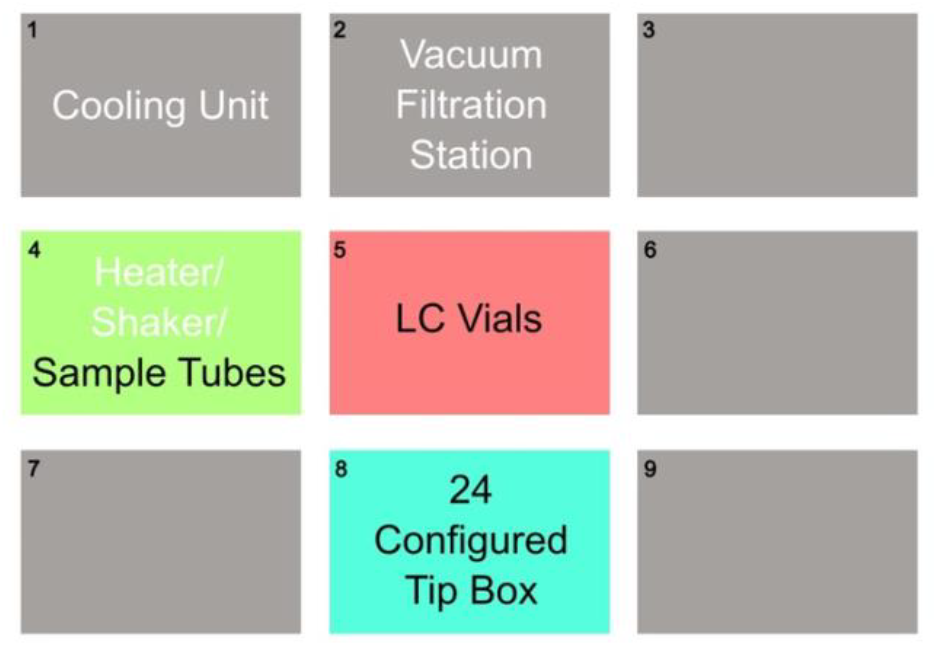
Map of the Bravo deck with labware configured for the ‘Transfer to LC Vials.pro’ protocol. Fixed labware (white text), 24-well microtube rack adapter with sample tubes (green), LC vial adapter rack (red), 24 tip box (turquoise), and positions with no movable labware (grey).

**Figure 6.**
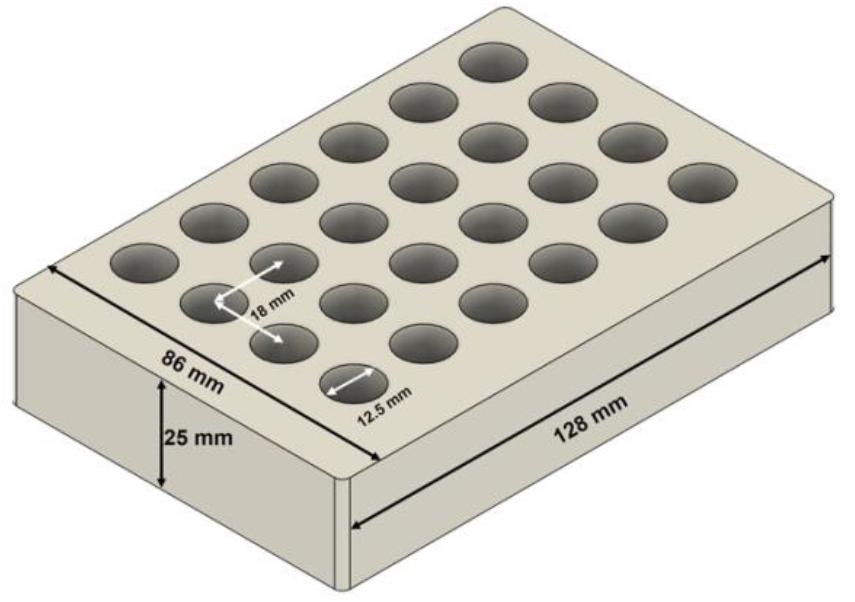
Dimensions of the 3D printed LC vial adapter rack.

#### Preparation of experiment-specific quality control (QC) pooled sample

1. Start the Agilent Bravo automated liquid handling platform, the heating shaking station, and the vacuum filtration station pump.
2. On the VWorks software, load the ‘QCPool.pro’ (access on GitHub). A prompt will appear to initialise the device now, if the device has not already been initialised. Click ‘Yes’ if applicable.
3. The form file ‘QCPool.VWForm’ (access on GitHub) will open automatically.
4. Retrieve metabolite extracts from ultra-low temperature freezer and centrifuge at 21,130 ×g for 5 min at 4°C.
5. Place the microtube rack adapter with microcentrifuge tubes containing samples (lids open) on position 5, a full 96 tip box on position 8, an empty tip box on position 9, and a microtube rack adapter with an open 1.5 mL microcentrifuge tube in A1 on position 6 of the Bravo deck (Figure 7).
6. Create a .csv file in Microsoft Excel to list the source sample plates and wells to be used in the protocol. The sheet should have two columns labelled ‘plate name’ (e.g. plate 1, plate 2…) and ‘sample well’ (e.g. A1, A2, A3…). Ensure that the file accurately replicates the position of the tubes in the source microtube adapter rack, then save the file and ensure that the data path on the form file leads to this sheet.
7. Select the desired volume of each sample to be transferred to the QC pool using the drop-down menu on the form file. Then click ‘Start’ on the form file to begin the protocol.
8. A dialogue box will appear with the message ‘Please place a new source plate on location 5’. Ensure that the source plate has been placed on position 5 and click ‘Continue’. A second dialogue box will appear, prompting the user to pause the protocol and update the tip state editor. Click ‘Pause and Diagnose’, then click the ‘Tip State Editor’ button in the form file. Ensure that the tip state editor matches what is in the boxes on positions 8 and 9 of the Bravo deck and that there are enough tips to transfer all of the samples. Then click ‘OK’ in the ‘Tip State Editor’ box, and click ‘Continue’ in the ‘Scheduler’ box.
9. The Bravo will then transfer the specified volume of each sample (one at a time) to the microcentrifuge tube in the microtube rack adapter on position 6 to generate the QC pool.
10. If more than one plate of samples has been listed on the excel input file, the protocol will loop back to step 8.

**Figure 7.**
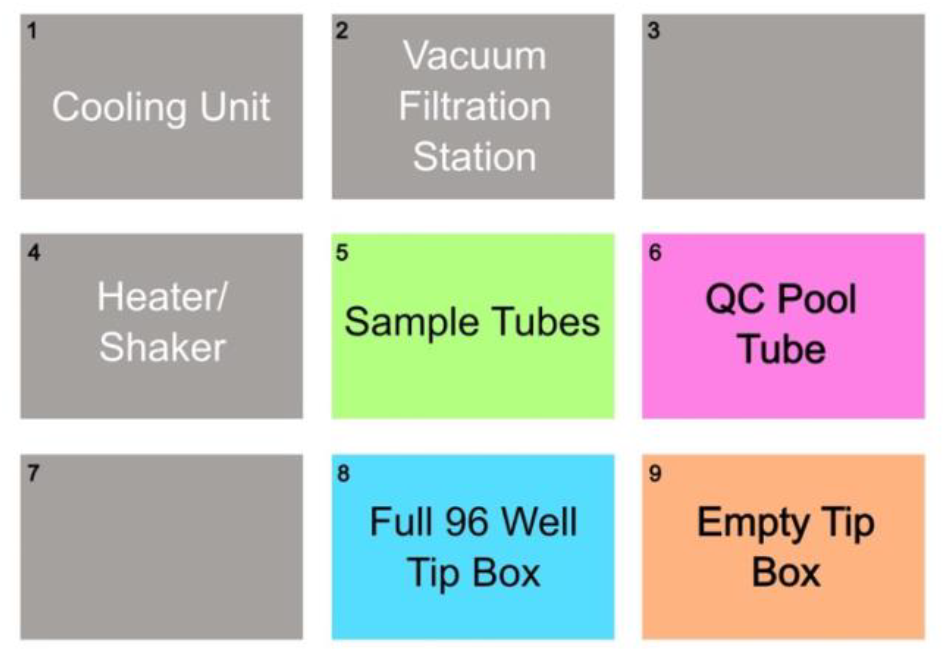
Map of the Bravo deck with labware configured for the ‘QCPool.pro’ protocol. Fixed labware (white text), 24-well microtube rack adapter with sample tubes (green), LC vial adapter rack (red), 24-well microtube rack adapter with the QC Pool tube (pink), full 96 well tip box (blue), empty tip box (orange), and positions with no movable labware (grey).

## Results and Discussion

To assess the automated metabolite extraction workflow, 192 Lysogeny Broth (nutritionally rich bacterial culture medium) samples were aliquoted and processed in two batches of 96 samples each using the Agilent Bravo ‘1in10 Metabolite Extraction_Liquid Sample_Microtubes.pro’ protocol. Each set of 96 metabolite extracts was then randomised and analysed by liquid chromatography – mass spectrometry (LC-MS) using the rapid HILIC-Z ion mobility mass spectrometry (RHIMMS) method (Pičmanová et al., 2022). The raw LC-MS data files were processed using the Agilent MassHunter software suite, and the obtained data was statistically analysed using MetaboAnalyst 6.0 (Pang et al., 2024). The data was log^10^-transformed, Pareto-scaled and visualised using principal component analysis (PCA) and partial least squares – discriminant analysis (PLS-DA).

The sample replicates bunched closely together and did not show differences between samples within a batch or across batches on the PCA (Figure 8a) and PLS-DA (Figure 8b) 2D scores plots, suggesting good technical replicability of the automated sample extraction workflow. We also looked at individual metabolite abundances across the 192 samples and observed similar peak intensities across the batches, two of these are plotted here (Figure 8c).

**Figure 8.**
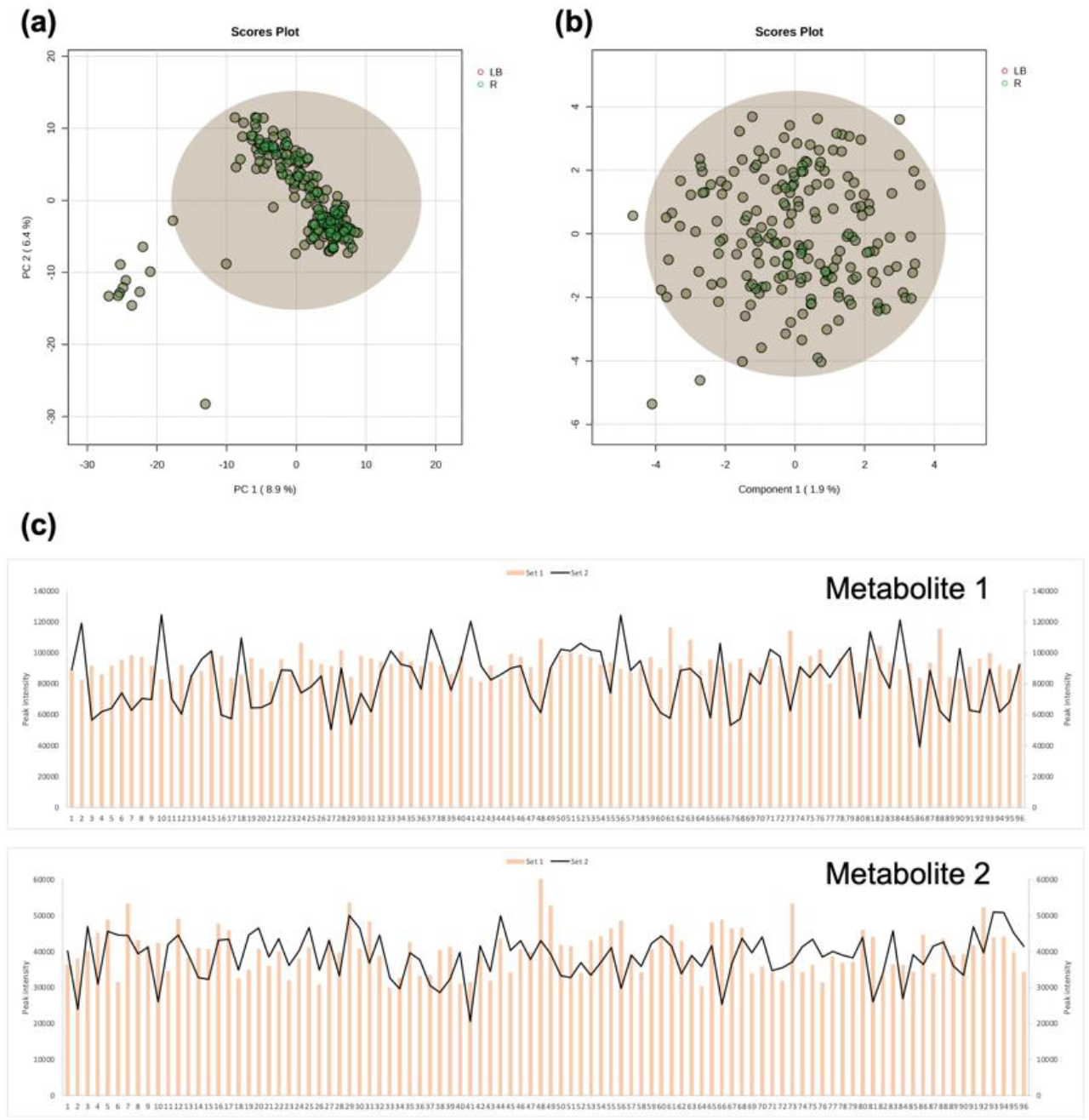
PCA (a) and PLS-DA (b) 2D scores plots of 192 LB metabolite extracts processed on the Agilent Bravo automated liquid handling platform and run on RHIMMS as two separate batches of 96 samples each show negligible variation between the individual extracts. (c) Bar and line plot of each batch of 96 samples, plotted for metabolites 1 and 2. The X-axis shows the 96 individual samples in each batch, and the Y-axes corresponds to peak intensity. The primary and secondary Y-axes have the same scale. One set of 96 samples is plotted as a bar graph (orange), and the other set of 96 samples as a line plot (black).

This automated sample aliquoting and metabolite extraction workflow is suitable for capturing the natural metabolite composition of complex biological mixtures/samples with high reproducibility. A total of 266 metabolites were consistently annotated in the dataset across the 192 samples. The direct comparison of metabolites in the individual batches, as well as across the two batches, suggests consistent extraction efficiency despite the inherent variation expected from the LC-MS methodology and post-run data processing workflows. The variability calculated as a percentage of the standard deviation from the mean was less than 8% per batch, with some of this variability being attributable to the LC-MS data acquisition and processing methods. Additionally, it is also worth noting that no sample position effects and individual metabolite extraction run effects that could potentially be introduced by repeated use of the Agilent Bravo workflows were apparent from the metabolite peak intensities.

Using this combination of workflows, the user is equipped with a robust, reproducible, and automated technique for metabolite extraction for global metabolomics analysis using LC-MS methods. The automated nature of this workflow frees up valuable user time, allowing, for instance, LC and MS instrumentation set-up.

We conclude that this degree of reproducibility is adequate to discover biologically relevant metabolic changes within complex biological samples. By taking advantage of the combination of high throughput and robustness afforded by the Agilent Bravo automated liquid handling platform, the preparation of hundreds of metabolite samples has now become feasible. We believe this application will be indispensable for large-scale engineering biology and clinical research projects.

## Author Contribution

FM, AM and TM conceived this project, designed and carried out the experiments, and wrote the manuscript. FM, AM, SP and MAC wrote the software files. All authors read and approved the manuscript.

## Conflict of Interest

The authors declare no competing financial interests.

## Notes

https://github.com/EdinOmics/Automated-metabolite-extraction-workflow-on-Agilent-Bravo-liquid-handling-platform/branches

### Competing Interest Statement

The authors have declared no competing interest.

